# Estimating null and potent modes of feedforward communication in a computational model of cortical activity

**DOI:** 10.1101/2021.10.18.464855

**Authors:** Jean-Philippe Thivierge, Artem Pilzak

## Abstract

Communication across anatomical areas of the brain is key to both sensory and motor processes. Dimensionality reduction approaches have shown that the covariation of activity across cortical areas follows well-delimited patterns. Some of these patterns fall within the “potent space” of neural interactions and generate downstream responses; other patterns fall within the “null space” and prevent the feedforward propagation of synaptic inputs. Despite growing evidence for the role of null space activity in visual processing as well as preparatory motor control, a mechanistic understanding of its neural origins is lacking. Here, we developed a mean-rate model that allowed for the systematic control of feedforward propagation by potent and null modes of interaction. In this model, altering the number of null modes led to no systematic changes in firing rates, correlations, or mean synaptic strengths across areas, making it difficult to characterize feedforward communication with common measures of functional connectivity. A novel measure termed the null ratio captured the proportion of null modes relayed from one area to another. Applied to simultaneous recordings of primate cortical areas V1 and V2 during image viewing, the null ratio revealed that feedforward interactions have a broad null space that may reflect properties of visual stimuli.

## 1 Introduction

Understanding how distinct areas of the brain communicate with each other is at the heart of neuroscience. Neurons in a given area receive synaptic afferents from thousands of upstream cells, and it remains unclear how this vast information is integrated to generate sensory, cognitive, and behavioral outcomes.

A number of measures aim to capture the interactions between neurons across brain areas, including functional connectivity [1, 2], Granger causality [3, 4], and information-theoretic measures such as transfer entropy [5–7]. With some notable exceptions including population coding [8], these measures typically produce estimates over all pairs of elements, resulting in prohibitively large matrices of interactions. With recent developments allowing for simultaneous recordings on the order of 10,000 neurons [9], these interactions would involve ~1,000,000 paired comparisons.

Several approaches have been proposed to reduce the sheer volume of neural data and summarize neuronal population activity using a handful of dimensions that reflect broad interactions within local circuits [10–15]. These analyses suggest that reliable patterns of covariation amongst cells, termed “neural modes", emerge from the collective behavior of large neuronal groups. These neural modes are reported in early visual cortex (V1), where the activity of large groups of cells is captured by linear models with only a few dimensions [15, 16].

Furthermore, feedforward communication between V1 neurons and downstream extrastriate area V2 [17] is described by a limited “neural space” whereby several of the activity patterns in V1 do not activate V2 targets. The presence of such neural space is not restricted to visual cortex, but is also reported in areas responsible for motor control [18–23] and is linked to movement preparation [18], learning [22], and working memory [24]. In the context of hand reaching tasks, the challenge is that cortical regions involved in the preparation of a movement are also involved in controlling the muscles responsible for initiating motor commands. A solution is that cortical activity related to the preparation of a movement occupies a null space where it does not generate a motor command. Upon movement initiation, neural activity switches to a potent space that communicates to the muscles involved in hand reaching [18]. In both visual and motor domains, neural activity can be described by “potent modes” that propagate their activity to downstream sites and “null modes” that yield no marked response.

Despite the ubiquity of null and potent modes across cortical areas, few computational models exist to capture how neuronal circuits control modes of neural activity in order to gate feedforward communication. Some models work by generating random vectors that are rotated along different axes to create null and potent modes [16, 18]. Unfortunately, these models offer no description of the underlying neural activity required to generate these modes. Other models are hard-wired to perform feedforward gating of neural activity [25] but offer no systematic way to control the transmission of null and potent modes. Finally, some models learn to generate low-dimensional representations of a signal [26, 27] but focus on activity within a single brain area.

Here, we developed a simplified mean-rate model that allowed us to systematically control the feedforward transmission of neural modes. The model revealed that different subsets of modes can be activated without notable changes in firing rates, correlations, or the mean strength of synapses, raising concerns for studies that aim to infer information transfer between brain areas via functional connectivity.

Going further, we created an unbiased measure, termed the null ratio, that captured the proportion of null modes relayed from one area to another. This measure improves upon previous estimates of null and potent modes [18] that yield an underestimation of effect sizes. We applied this measure to simultaneous multielectrode recordings in areas V1 and V2 of primate visual cortex. During a passive image viewing task, interareal communication was dominated by a high proportion of null modes and a comparatively smaller number of potent modes, potentially capturing the statistics of sensory input presented during the task.

## 2 Results

### 2.1 Mean-rate model of null and potent neuronal interactions

We began with simulations where a mean-rate “sender” network received a frozen Gaussian signal and selectively propagated its activity in a feedforward fashion to a “receiver” area (Fig. 1a). Synaptic weights **W**_0_ between the two areas were adjusted such that a specific set of neural modes were set to null. This was achieved by solving a linear equation that projects the activity from the sender network to the receiver network (see Materials and Methods, equations (6–16)). Unless otherwise stated, 10 seconds of simulated activity was generated for each run of the model.

**Figure 1.**
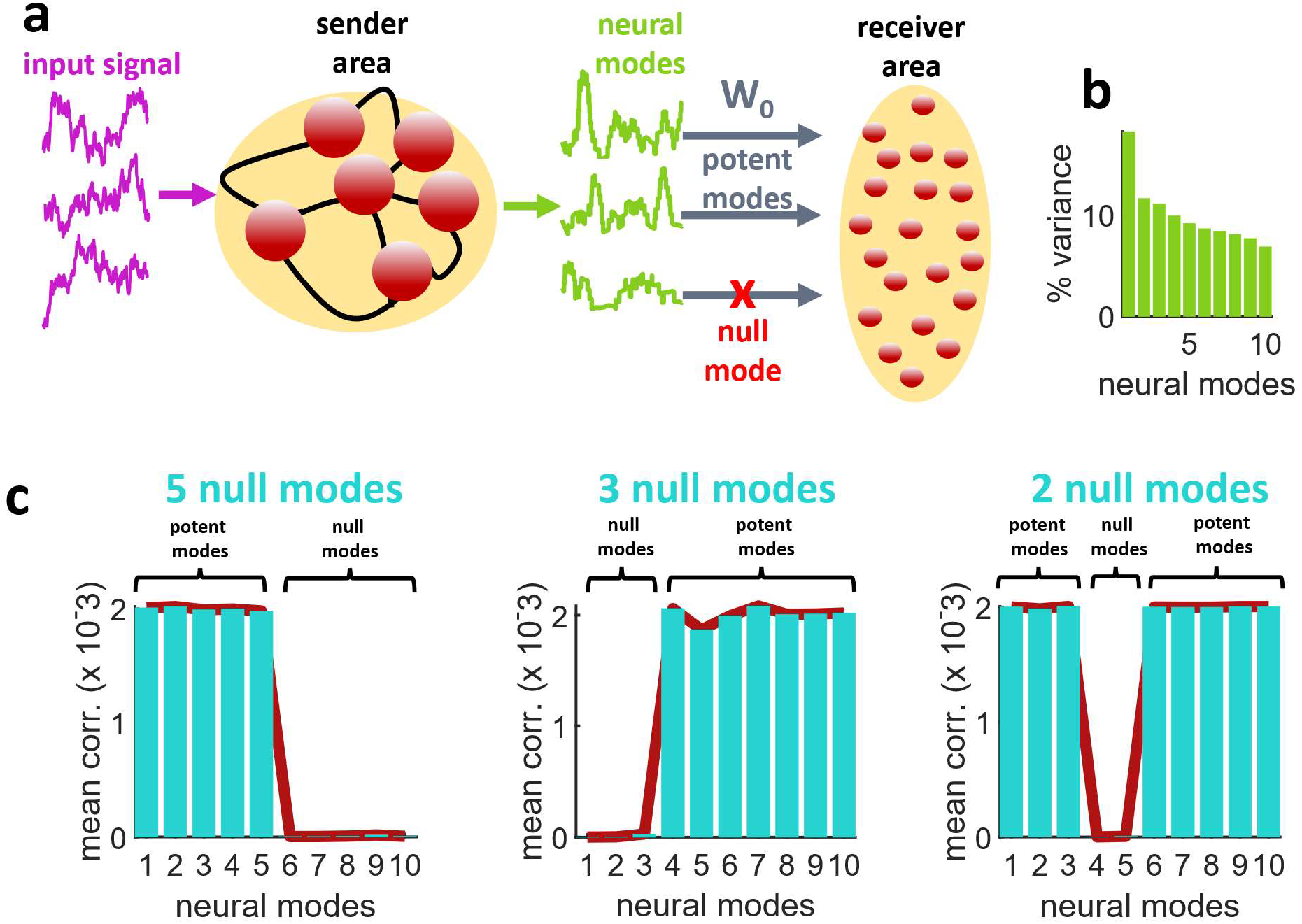
A model of feedforward communication between two neuronal circuits. (**a**) Simplified model where a recurrently-connected sender area receives an input signal and projects to a receiver area via a set of connection weights (W0) where potent neural modes are transmitted forward while null modes are not. (**b**) Percentage of variance explained by neural modes (*N*=10). (**c**) Across different simulations that altered the number and origin of transmitted neural modes, potent modes yielded a higher correlation between sender and receiver areas than null modes (*N*=10). Results shown were obtained from a receiver area with (solid lines) and without (vertical bars) lateral connectivity.

The presence of mode-specific interactions between the sender and receiver areas was readily observed by extracting neural modes of each area, respectively (see Materials and Methods, equations (7–8)). Modes were sorted by rank such that the first mode explained the highest proportion of variance (Fig. 1b). Three examples with different sets of null modes are shown in Fig. 1c. Given a full-rank singular value decomposition, the total number of modes is equal to *N* (the number of neurons in each area of the model). Similar results were obtained regardless of the inclusion of lateral connections between the receiver neurons (equation (16)). Even though correlations between modes of the two areas were low, stronger positive correlations emerged between potent modes, and weaker correlations between null modes. In fact, in a noiseless scenario, correlations between null modes would reach zero. The ability of the model to turn off particular null modes was possible for modes that explained a low proportion of variance (Fig. 1c, left panel) as well as modes that explained a higher proportion (Fig. 1c, middle and left panels).

To illustrate the impact of null and potent modes on the receiver area, a simulation was performed where either a single null or potent mode was activated. A simple example with three neurons in each of the sender and receiver areas is shown in Fig. 2a. The mode capturing the largest proportion of variance was set to null, while the remaining modes were potent. Activating only the null mode resulted in activity in the sender but not the receiver area. Conversely, activating a potent mode resulted in a response across both areas. Importantly, the overall distribution of activity in the sender area remained similar for both null and potent modes (Fig. 2b), showing that results were not due to a trivial attenuation of activity when generating null activity in the sender area.

**Figure 2.**
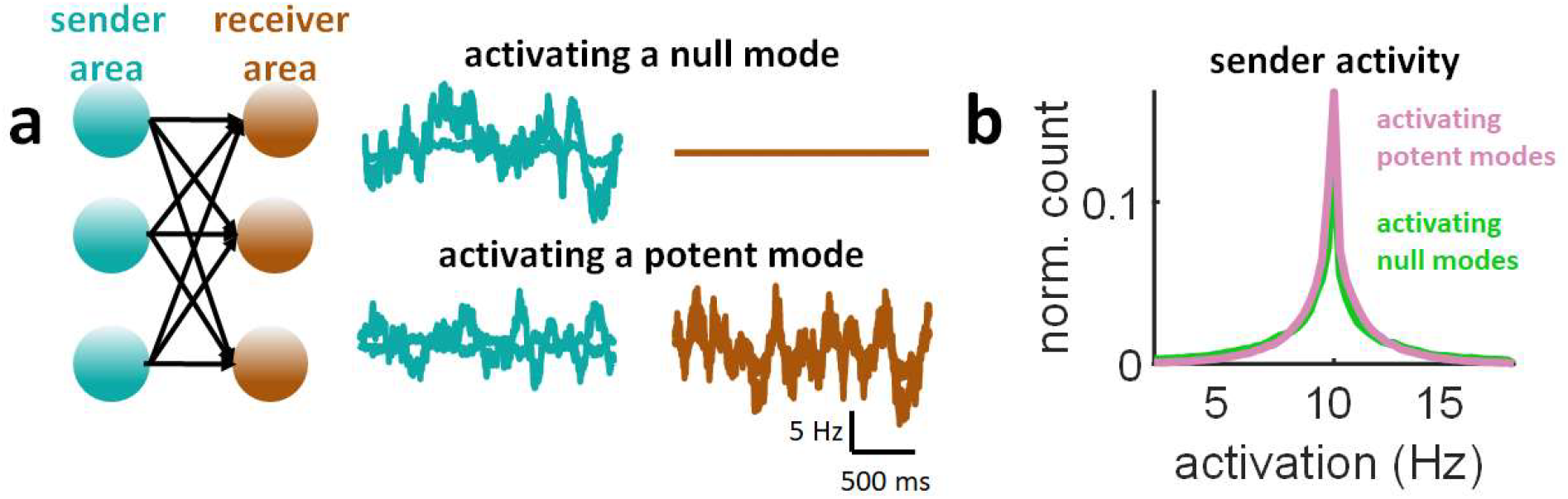
Toy example of a network with three sender and receiver neurons. (**a**) Activating a null mode led to no response in the receiver area. Conversely, activating a potent mode generated a response in both the sender and receiver areas. (**b**) Distribution of activity from the sender area when activating either a null or potent mode.

In a larger simulation, the outgoing connectivity of *N*=100 sender units was configured to propagate *N*/2 potent modes in a feedforward fashion. Mean pairwise correlations between modes of the sender and receiver areas reflected null and potent modes (Fig. 3a, left panel). A similar result was obtained when increasing the size of the population (*N*=500 with 400 potent modes and *N*=800 with 700 potent modes) (Fig. 3a, middle and right panels). In turn, synaptic weights between the sender and receiver areas yielded a Gaussian-like distribution centered near zero, with no clear structure (Fig. 3b).

**Figure 3.**
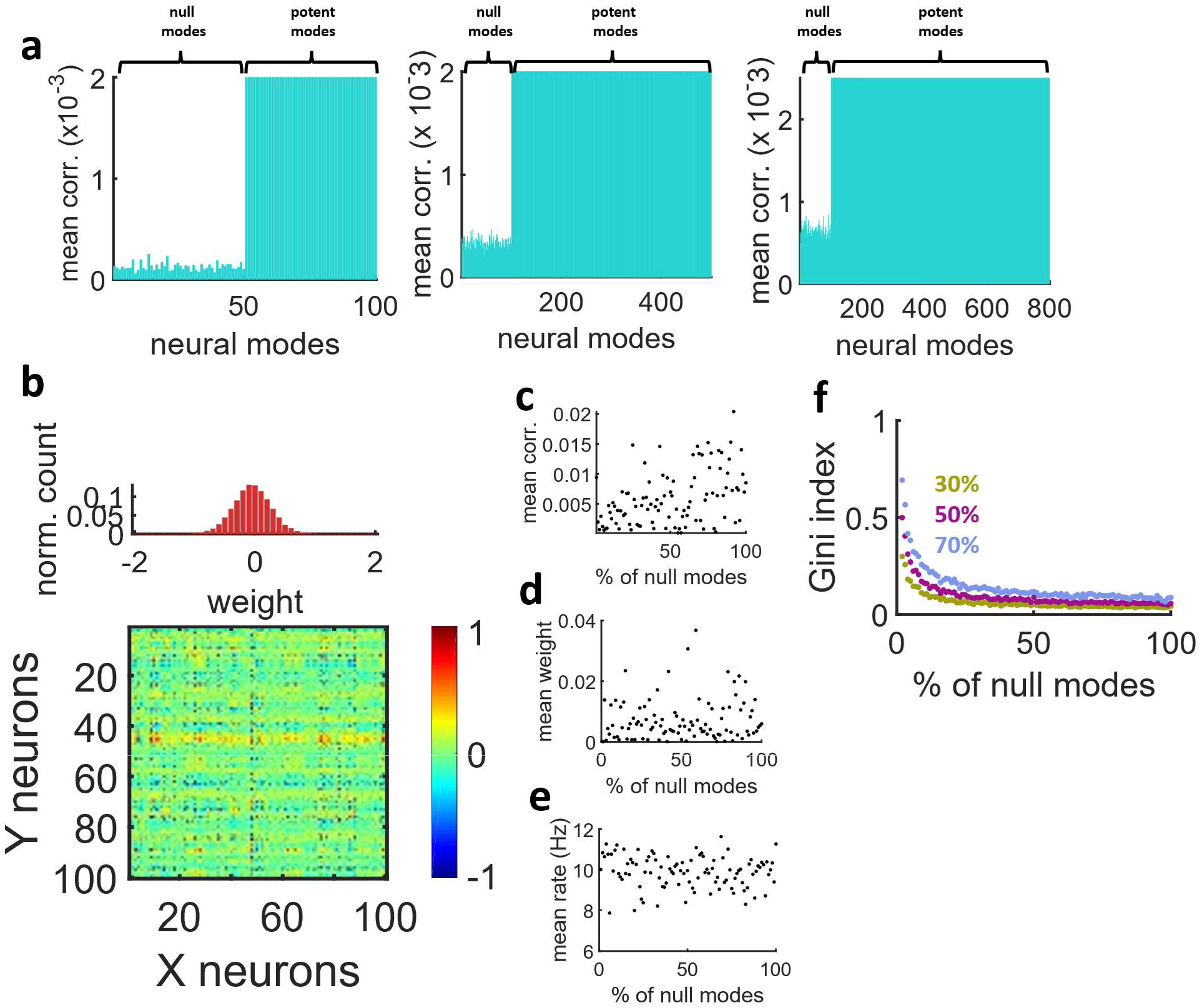
Null modes were not straightforwardly reflected in synaptic weight distribution, firing rates, pairwise correlations, or mean synaptic weights. (**a**) Correlation between modes of the sender and receiver areas with *N*=100, 500, or 800 units and 50, 400, and 700 potent modes, respectively. (**b**) Connection weights between sender and receiver areas. (**c**) Absolute mean pairwise correlation in different runs of the model where the percentage of null modes was altered. (**d**) Absolute mean synaptic weight in relation to the percentage of null modes. (**e**) Mean activity of the sender area versus null modes. (**f**) Gini index of the feedforward connectivity matrix. A threshold was applied to convert real-valued weights to a binary adjancency matrix where a given percentage of strongest connections were preserved.

Thus, the propagation of potent and null modes was not apparent from basic features of synaptic connectivity. Further, configuring a network with a combination of null and potent modes did not yield a trivially sparse matrix where null modes were readily identifiable.

A number of independent runs of the model examined a broad range of null modes. In these simulations, the proportion of null modes showed no relation to either mean absolute pairwise correlations across the two areas (Fig. 3c), mean absolute synaptic weights (Fig. 3d), or combined mean firing rates of the sender and receiver areas (Fig. 3e).

A measure termed the Gini index [28] was employed to assess the impact of null modes on network sparsity. First, synaptic weights were converted to a binary adjacency matrix by a threshold that retained the highest 30%, 50%, or 70% of weights. Then, the Gini index was calculated across a range of null modes (Fig. 3f). While the Gini index diminished between 0-50% of null modes, the change in Gini index between 50-100% of null modes was small.

Hence, it was difficult to directly observe null modes from basic features of neural activity or functional connectivity in the model, including correlations, weights, firing rates, and network sparsity. While the correlation between individual modes of the sender and receiver areas was modulated by setting particular modes to null, the overall correlation between the activity of the two areas remained largely unaffected.

Results of the rate-based model were extended to a scenario where the sender network was comprised of *N* =500 spiking neurons (see Materials and Methods, equations (17–19)) (Fig. 4a). Ten seconds of activity were generated with a set proportion of null modes. The receiver network followed a bistable regime with sharp transitions between a low and higher state of activity. The correlation between modes of the sender and receiver areas was low for null modes, and higher for potent modes (Fig. 4b). Thus, the proposed model allowed for a systematic control of null and potent modes propagated from one area to another.

**Figure 4.**
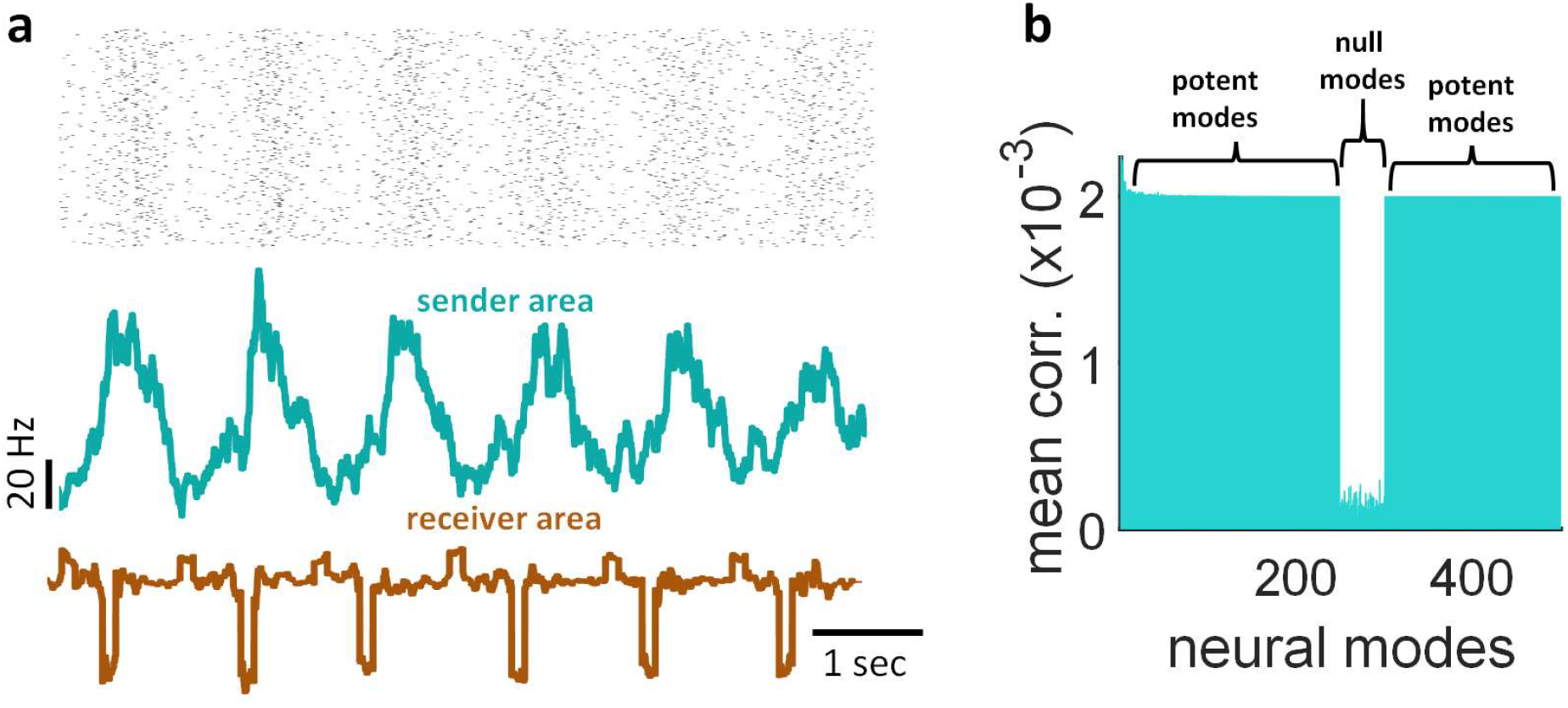
Propagating null and potent modes of neural activity in a spiking network. (**a**) Top: spike raster from a sender network (*N*=500 units) where a subset of modes (250 to 350 in rank) were set to null. Bottom: sum of population activity over time. (**b**) Correlation between modes of the sender and receiver areas.

### 2.2 Measuring null space communication

To measure the null space of communication between two neural areas, we began by describing the propagation of activity from a sender area **X** to a receiver area **Y** as a weighted sum, **Y** = **W**_0_**X** + ***c***, where ***c*** is a constant term and **W**_0_**X** is a projection of activity **X** onto a set of weights **W**_0_ that reflect the influence of the sender region on the receiver network (see Materials and Methods). Potent and null modes of propagation were obtained by the row-space 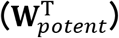 and null space 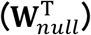 of **W**_0_, respectively [18]. The transmission of potent and null modes from the sender to the receiver network was described by projecting the eigenvectors of **X** (denoted **V**) onto the row-space and null space of **W**_0_,

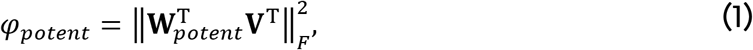

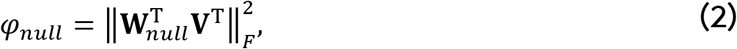

where 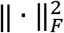 is the Frobenius norm. This norm was employed for ease of interpretation, as it is closely related to the variance of the expression inside the brackets. That is, subtracting the mean of each row before taking the norm yields the variance. The resulting expressions *φ_poten_* and *φ_null_* reflect the influence of the sender area along the potent and null space, respectively. Crucially, these measures require that the sender and receiver networks have the same number of neurons (*N*) in order to be meaningful. Otherwise, “funnelling” or “expanding” activity from the sender to the receiver areas would cause a spurious number of null or potent modes to emerge.

Equations (1–2) were computed across 10 simulations where the proportion of null modes was varied between 0-100%. In these simulations, the potent and null space of **W**_0_ crossed at a point where half of the neural modes (*N*/2) were null (Fig. 5a). A ratio of null and potent spaces was computed to obtain the proportion of null modes,

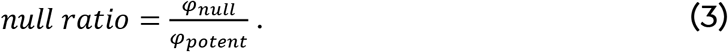

**Figure 5.**
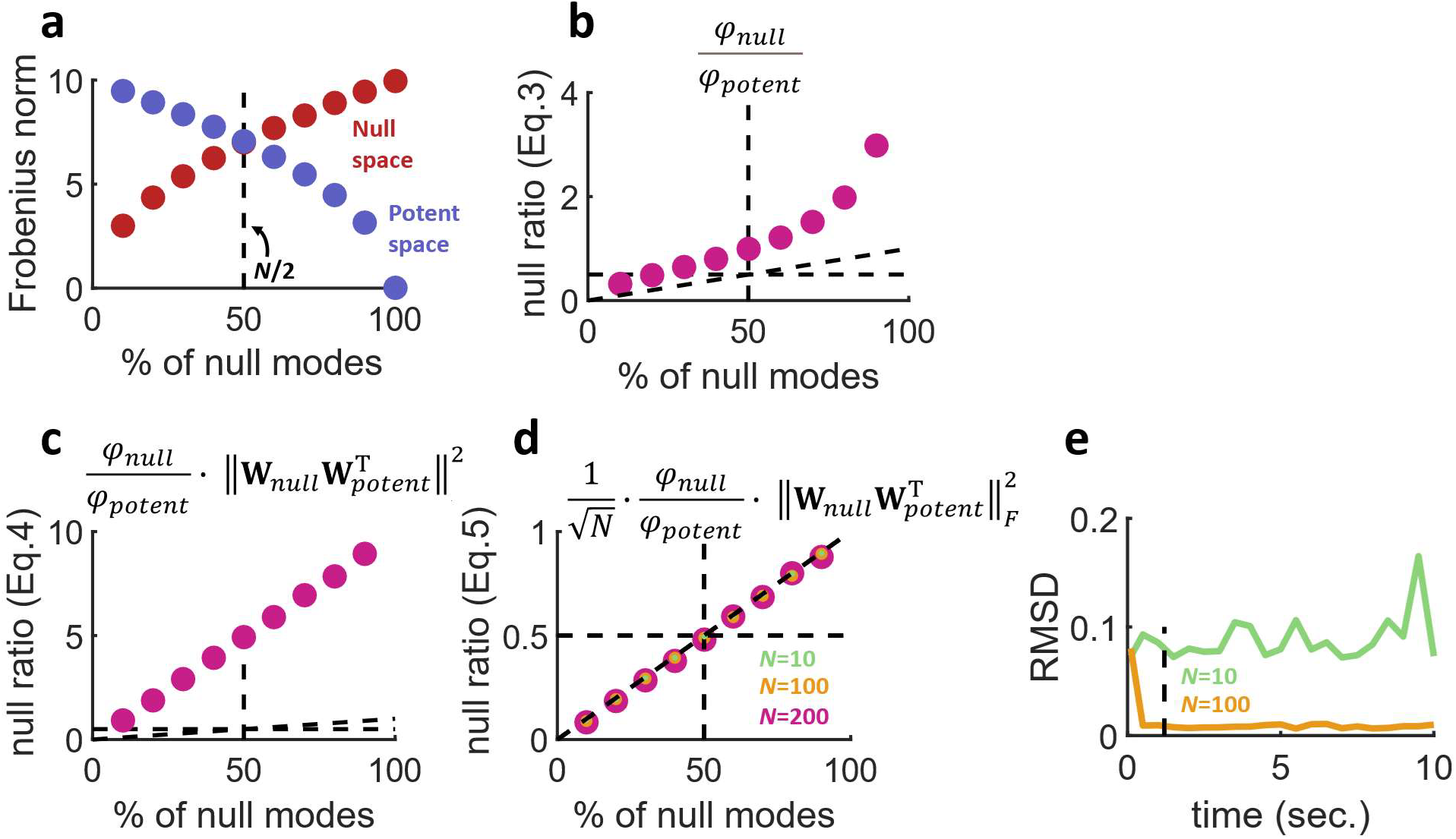
Estimating the null and potent space of neural communication. (**a**) The Frobenius norm of the null and potent space of connections W0 intersected at a location where approximately half of all modes are null (*N* =100). (**b**) The ratio of null and potent neural modes (equation (3)) provided an estimation of the proportion of null modes in the model, but also generated unwanted nonlinearity. (**c**) Correcting nonlinearity by adding a term to the ratio (equation (4)) led to an overshoot in prediction. (**d**) The overshoot in prediction was corrected by scaling the estimation with a factor inversely proportional to the square root of *N* (equation (5)). (**e**) Root mean square difference (RMSD) between actual and estimated null ratios across different durations of simulation. Vertical dashed line: one second of activity.

This measure, however, featured a non-linearity that overestimated the null modes (Fig. 5b) [18]. This non-linearity can be corrected by adding a term to equation (3),

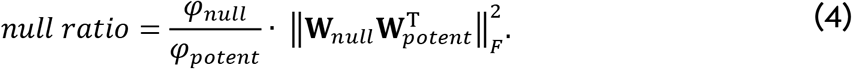

This new estimator was linearly related to the null modes, but was still prone to a large overestimation bias (Fig. 5c). This bias was corrected by scaling equation (4) by 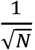, canceling out the squared norm of equations (1–2),

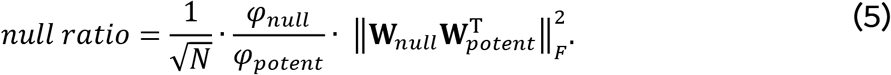

This final estimator correctly reported the proportion of null modes independently of *N* (Fig. 5d). To determine how many data points were required to perform an adequate estimate of the null ratio, different runs of the model were performed where the number of simulated time-steps was altered. With 10 seconds of activity and *N*=100 neurons, the root mean squared difference (RMSD) between the actual and estimated null ratios was low. However, this difference increased when the time segment of activity was shortened (Fig. 5e). Thus, the estimated null ratio improved with longer segments of data. A low number of neurons (*N*=10) yielded a high error regardless of the time segment. With *N*=100, at least one second of activity (Fig. 5e, dashed vertical line) was required to provide a reliable estimate of null ratio.

In practice, the estimation of null modes may be applied to experimental data where the activity of a sender and receiver area is recorded simultaneously. An example is shown in the next section, based on cortical activity from visual areas V1 and V2.

### 2.3 Null and potent dimensions in cortical activity

Single-trial activity was analyzed where anesthetized primates viewed sinusoidal gratings and activity was recorded simultaneously in V1 and V2 areas (see Materials and Methods). These data focused on neurons from the superficial layers of V1 that project to middle layers in V2 [16]. These local projections constitute ~95% of afferents to V2 [29], ruling out a strong influence of surrounding areas. V1 activity was split into two groups of an equal number of neurons matching the number in V2. More precisely, a total of 111 neurons were recorded from V1 and 37 from V2. Two random samples of 37 neurons each were extracted from V1. The first sample was compared to V2, yielding “V1-V2” comparisons, while “V1-V1” comparisons were obtained by comparing the two V1 samples (Fig. 6a). The same approach was employed in related work [16]. The peri-stimulus time histogram of single-trial activity was regressed using the Matlab function *regress*, which performs multiple linear regression using the least-squares method [30] and returns the residuals. The activity of individual neurons was smoothed by averaging their firing rate over time using a sliding window of 100 ms.

**Figure 6.**
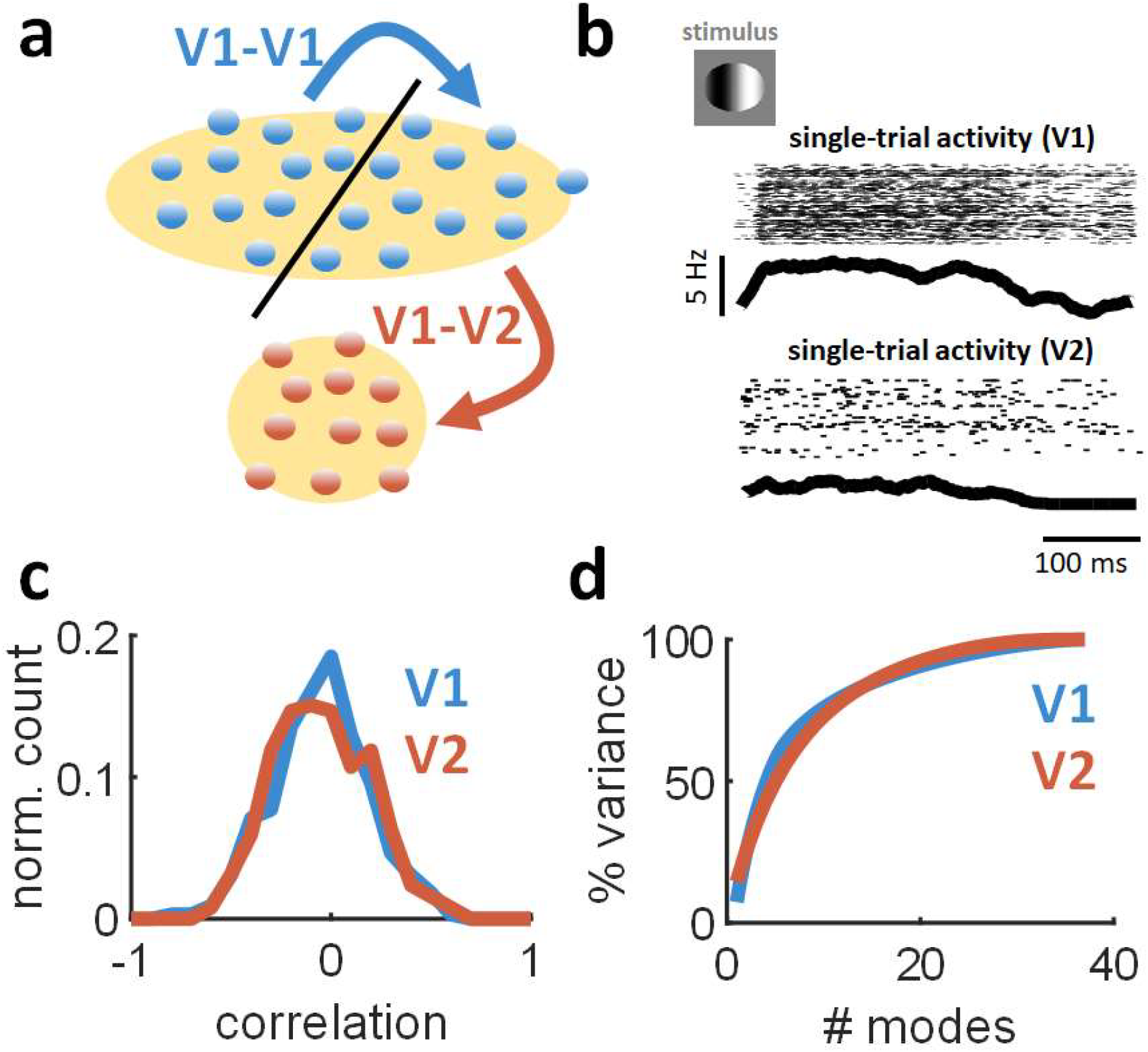
Analysis of feedforward interactions in cortical areas V1 and V2. (**a**) Simultaneously recorded V1 neurons were split into two even groups whose size matched the number of units recorded in V2. This arrangement of the data allowed for analyses of both V1-V1 and V1-V2 interactions. (**b**) Example of single-trial activity in simultaneous recordings of V1 and V2 neurons in response to oriented stimuli. Top: spike raster across all neurons. Bottom: summed population activity (smoothed using a 100 ms rolling window). (**c**) Distribution of pairwise correlations within cells of V1 and V2, respectively. (**d**) Variance explained by neural modes of V1 and V2 activity.

Single-trial activity was characterized by the activation of a large subset of V1 neurons with a rapid rise time (~50 ms) and slower decay (~150 ms) (Fig. 6b). A similar, albeit sparser, response profile was observed in V2. Intra-areal pairwise correlations were comparable between V1 and V2 [16] (p=0.28658, one-sided Wilcoxon rank sum test, n=28,800) (Fig. 6c).

Singular value decomposition was employed to examine the dimensionality of V1 and V2 activity (Materials and Methods, equations (7–8)). The variance explained by this analysis increased rapidly with the number of modes considered (Fig. 6d). The cumulative distribution of both V1 and V2 modes reached 100% of explained variance as the singular value decomposition approached full rank. A Wilcoxon rank sum test compared the cumulative distributions of V1 and V2 modes. This analysis revealed that the variance explained by V2 modes was lower than V1 (p=4.0535×10^−9^, one-sided Wilcoxon rank sum test, n=148). Hence, V2 activity was characterized by higher dimensionality than V1, in line with previous work [16].

Next, a linear mapping of activity from the sender area (V1) to the receiver area (V1 or V2) was obtained by ridge regression (see Materials and Methods). An example of V2 approximation based on smoothed V1 activity from a single trial is shown in Fig. 7a. The fit between observed and approximated activity was high (Fig. 7b): over all trials, the mean variance explained by ridge regression was 87.56% for V1-V1 interactions and 96.21% for V1-V2. Thus, while cortical neurons likely perform non-linear operations on their inputs [31], a majority of the variance in the receiver area was explained by a linearly weighted fit of V1 activity.

**Figure 7.**
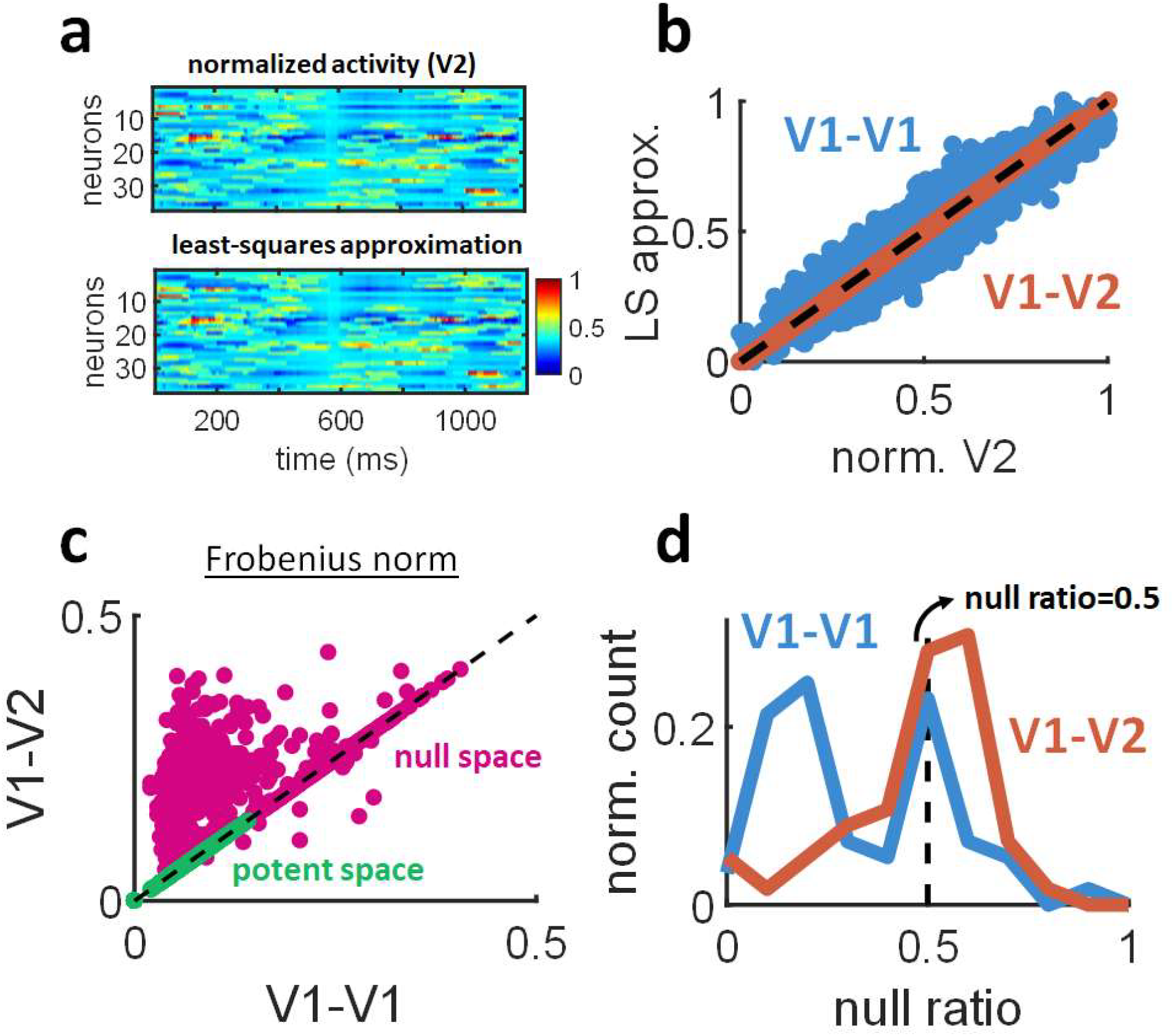
Null and potent modes estimated from simultaneous recordings of V1 and V2 activity. (**a**) Example of a single trial where V2 population activity was predicted by ridge regression of smoothed V1 activity recorded during the same trial. (**b**) Relation between V2 activity from panel (a) and its approximation by ridge regression. Individual circles are mean firing rates taken over 100 ms windows. Dashed line: unity. (**c**) Null and potent modes of V1-V1 and V1-V2 interactions. Individual circles are single trials. Each dot represents a value obtained from equations (1–2). (**d**) Distribution of null ratios. Dashed line indicates a null ratio of 0.5, corresponding to an equivalent number of null and potent modes.

Regression weights from V1-V1 and V1-V2 interactions were decomposed into null and potent modes by applying equations (1–2) across single trials. While the potent space was similar between V1-V1 and V1-V2 (p=0.999, one-sided Wilcoxon rank sum test, n=1,600), the null space of V1-V2 was markedly larger (p=4.4655e-06, one-sided Wilcoxon rank sum test, n=1600) (Fig. 7c). These results are in line with recent work showing that V1-V2 communication requires far fewer modes than the total available neural space [16].

To further compare null and potent modes, a bootstrap procedure was applied as follows. For each of 100 bootstrap steps, a random subset of neurons was extracted from V1 (matching the number of neurons in V2). Across 100% of these steps, the mean Frobenius norm (equations (1–2)) of the null space exceeded the potent space. Hence, V1-V2 modes were robustly characterized by a large null space.

Applying the null ratio (equation (5)) revealed that greater than half of V1-V2 modes fell within the null space (null ratio > 0.5) (Fig. 7d). By comparison, V1-V1 interactions yielded a bimodal distribution where a large proportion of modes fell in the potent space (null ratio < 0.5). The larger null space of V1-V2 interactions is consistent with findings of “private” V1 modes that have little predictive value on V2 activity [16]. This result cannot be accounted for by the number of neurons analysed (which was kept constant between V1 and V2) or pairwise correlations (Fig. 6c). Further, V2 dimensionality was greater than V1 (Fig. 6d), running counter to an explanation whereby a large number of null modes may be due to a lower number of V2 dimensions. Finally, 100 independent simulations of the mean-rate model were performed where the proportion of null modes was altered. These simulations showed that the null space can be manipulated without altering correlations (Fig. 3c), mean synaptic weights (Fig. 3d), or firing rates (Fig. 3e), suggesting that these factors may not have a substantial impact on estimating null and potent modes. Below, we discuss alternative explanations to the vast null space of V1-V2 interactions.

## 3 Discussion

This work described a mean-rate model of neuronal activity that enabled the control of null and potent feedforward modes of interaction between two brain areas. Based on this model, a novel, unbiased measure was developed to estimate the proportion of null modes. This measure was applied to simultaneous recordings of V1-V2 activity during stimulus viewing and revealed that most dimensions fell within the null space of feedforward communication between the two areas. The proposed measure of null ratio is model-independent and may be broadly applicable to other brain areas including motor cortex [18], hippocampal-entorhinal cortex [32], cortico-basal-ganglia circuits [33], and thalamocortical connections [34] under the condition that a sufficient number of neurons and time-steps are available to generate a reliable estimate (Fig. 5e).

### 3.1 Functional roles of null and potent modes

The ability of neural circuits to route feedforward activity through null and potent modes has implications that extend to sensory, motor, and cognitive domains. In sensory regions, dynamically activating particular modes may allow information to be flexibly routed from primary areas to higher cortical centers that perform multimodal integration [35]. In this way, neural circuits that control the propagation of null and potent modes may shape the integration and segregation of activity across regions [36]. Integration across regions may be achieved by the activation of potent modes, while segregation would arise through null modes. Speculatively, potent modes may also provide a neural basis for selective attention [37], whereas unattended sensory input may remain in the null space.

In brain regions responsible for motor control, null and potent modes mediate the preparation and execution of task-related movements [18]. During the preparation stages of a motor command, motor cortex activity resides in the null space, thus preventing movement execution. At the offset of the preparation stage, activity switches to potent modes that drive movement in accord with an appropriate motor plan. Thus, the rapid coordination of null and potent modes allows motor actions to be adequately planned and carried out in cortex.

### 3.2 Origins of the large null ratio in V1-V2 communication

What may explain the large null space of V1-V2 interactions (Fig. 7d)? One contributing factor may be the brain state of animals during experimental recordings. The use of anesthesia induces global, low-dimensional fluctuations across cortical regions [38]. These fluctuations may contribute to the scope of null space interactions across visual areas. However, a limitation of this explanation is that anesthesia-induced fluctuations are typically one-dimensional [15]. By comparison, our results suggest the presence of several modes in the null space of V1-V2 communication (Fig. 7d). Hence, the low-dimensional null space induced by anesthesia would not be adequate to explain the large null space found between V1 and V2.

Alternatively, the large null space of V1-V2 communication may be explained by the few dimensions required to encode the artificial stimuli employed experimentally. Indeed, only a few PCA dimensions would be required to encode oriented gratings compared to natural images, which typically require a dozen or more dimensions to capture most of their variance [39]. Because subjects were presented with gratings that required only a few modes to encode, the neural space of V1-V2 communication may encompass a broad null space without affecting sensory processing. Consistent with this explanation, Semedo et al. [16] found that when subjects were presented with natural movies of increasing duration, a greater number of neural dimensions was required to account for V1-V2 interactions, thus reflecting increased coding requirements. Further experiments that directly compare simpler and more complex stimuli will be required to further validate this proposal.

### 3.3 Implications of the proposed model

Our results challenge two fundamental assumptions about neuronal communication between brain areas. Firstly, a widespread assumption concerns the neural origin of sensory gating. Broadly speaking, gating is defined as the selective modulation of cortical inputs. Gating is generally assumed to be performed at the target site, for instance at the output of the cortical motor system [40]. However, our model suggests that gating may be controlled by the pattern of synaptic connections between sender and receiver areas of cortex. This opens the possibility that gating may be dynamically controlled and subject to short- and long-term synaptic plasticity [14].

A second assumption regarding neural communication that is called into question by our results is that the propagation of null and potent modes emerges from a balance of excitation and inhibition [25, 41]. According to this explanation, null modes correspond to states of detailed balance where excitatory and inhibitory inputs cancel out at the target site. Conversely, balance-breaking activity would form potent modes of transmission. In contrast with this explanation, the mean-rate model showed that null modes can emerge without requiring presynaptic activity to cancel out (Fig. 2a). In the model, null modes were not caused by anti-correlated activity in the sender area. Rather, null modes were controlled by the precise configuration of synaptic weights between sender and receiver areas.

A key contribution of the mean-rate model is that it highlights limitations of functional and structural connectivity in probing the interactions between brain areas. Specifically, simulations showed that pairwise correlations are a poor predictor of null space. Drastically altering the size of the null space had no systematic impact on mean pairwise correlations between the two neural areas (Fig. 3c). The implication of this finding is that it may be possible for neural activity to yield large correlations despite most modes falling within the null space. Conversely, low correlations could be obtained from activity where a majority of modes are potent. Hence, functional connectivity may offer misleading indications of the communication bandwidth between brain areas.

Furthermore, synaptic connectivity (Fig. 3d) and mean firing rates (Fig. 3e) were poor indicators of the breath of null space interactions. Thus, the need for adequate measures of null modes (equation (5)) may not be circumvented by common network statistics [1].

### 3.4 Future work and conclusions

One factor not examined here in the mean-rate model is the dimensionality of neural modes within the recurrent network formed by sender neurons. While this consideration has been the subject of extensive theoretical work [26, 41–44], the focus of the current work was the feedforward propagation of neural modes, and not their origin within recurrent circuits. Further work that examines both aspects of communication in a unified framework would provide an increased understanding of how interactions both within and across brain areas give rise to sensory perception and motor planning.

In conclusion, this work combined computational modeling and experimental analyses in order to describe avenues by which neural circuits may control feedforward communication across brain areas. A novel measure termed the null ratio was provided to account for null features of neural activity that do not propagate across areas. Applied to neural interactions in early visual cortex, this measure revealed that a large portion of V1-V2 activity fell within the null space. These results open the door to further applications of the null ratio across sensory and motor systems, linking inter-areal interactions to both cognitive and behavioral processes.

## 4 Materials and Methods

### 4.1 Mean-rate model

The model considered two brain areas of *N*=100 neurons each (unless otherwise stated), communicating via feedforward synaptic connections (Fig. 1a). Focusing on linear dynamics, activity **X** ∈ *ℜ^N×T^* for time-steps *t*, …, *T* in the sender area was modeled by a neural integrator [34–36],

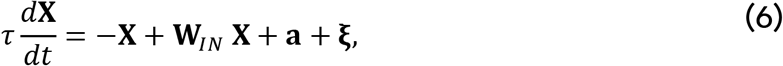

where internal connection weights **W**_*IN*_ were drawn from a Gaussian distribution 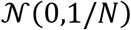 without self-connections, **ξ** is a frozen Gaussian input signal drawn from 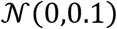, *t* is a time-step, *a* is a tonic input set to 10 Hz, and *τ*=10 ms is an integration time constant. The activity of each unit was smoothed using a rolling window of 100 ms to mimic the processing of experimental data as detailed below.

Next, we obtained the singular value decomposition of **X**,

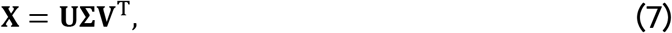

and computed neural modes by projecting the neural activity onto eigenvectors **V**,

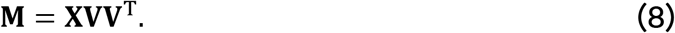

These modes reflect time-dependent signals along individual dimensions of **X** [48]. Activity from the sender area was assumed to propagate to the receiver area (**Y**) via a set of weighted feedforward connections **Z** ∈ ℜ^*N*×*N*^, defined as a random matrix with independent Gaussian elements 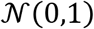. The resulting activity in the receiver area was thus

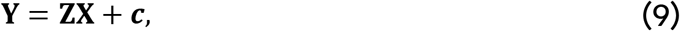

where ***c***=10 Hz is a constant bias. However, instead of **Y** being impacted by **X** directly, we sought for **Y** to be influenced by neural modes **M** (equation (8)), hence

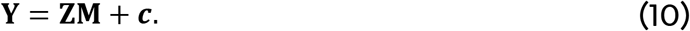

By expanding **M**,

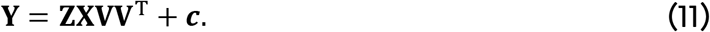

Going further, we would like **Y** to be influenced by some, but not all, of the neural modes. For this purpose, equation (11) was rewritten as

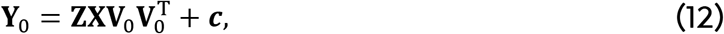

where **Y**_0_ describes the activity of the receiver area in response to null and potent modes transmitted by the sender area. The matrix **V**_0_ is the same as **V** (equation (7)), but with certain columns, corresponding to null modes, set to zero. We assumed a set of weights **W**_0_ that project activity from the sender to the receiver area. These weights should be distinguished from **V**_0_, the matrix containing eigenvectors of the sender area. Activity from the sender area **Y**_0_ can be substituted for **W**_0_**X** + ***c*** in equation (12). Then, **W**_0_ is sought such that

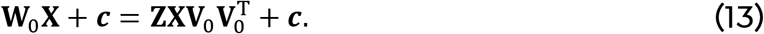

This is found by

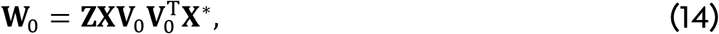

where “*” is the Moore-Penrose matrix inverse, employed here because **X** is not a square matrix given that there are typically more time points than neurons. Finally, the activity of the receiver area when receiving null and potent modes from **W**_0_ is obtained by

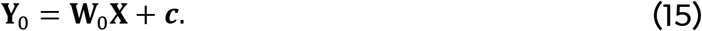

In a scenario that included lateral connectivity amongst receiver neurons, the above equation became

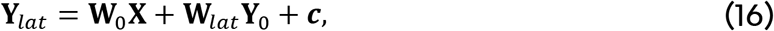

where **Y**_*lat*_ is the activity of the receiver units and **W**_*lat*_ are lateral connections between these units. These connections were i.i.d. elements drawn from a Gaussian distribution and matched the range of values in **W**_0_ (Fig. 3b).

Importantly, equation (14) should not be interpreted as a biological learning rule, but rather as a way of computationally generating feedforward connection weights whose structure allows for the systematic control of null and potent modes.

### 4.2 Spiking network model

Simulations were performed where the sender network was composed of *N* = 500 recurrently-connected spiking neurons [39, 49]. The membrane potential of individual neurons evolved according to

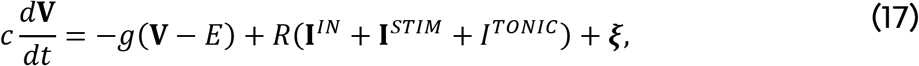

where *g*=0.01 pS is a leak conductance, *E*=−65 mV is a reversal potential, *R* =10 Ω ∙ cm is the membrane resistance, *I*^*TONIC*^ =0.5 nA is a constant current, **I**^*STIM*^ is a current induced by the stimulus (modeled by frozen Gaussian noise with 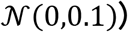), and 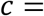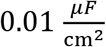 is the membrane capacitance. The intrinsic current 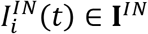,

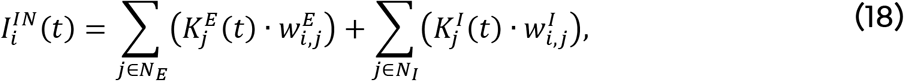

describes the contribution of the surrounding network activity to an individual unit *i* at time *t*, where 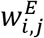 is the outgoing connection strength from one excitatory (E) neuron *i* to a neuron *j*, 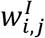 is the outgoing strength from an inhibitory (I) neuron, *N_E_* and *N_I_* are the total numbers of excitatory and inhibitory neurons, and

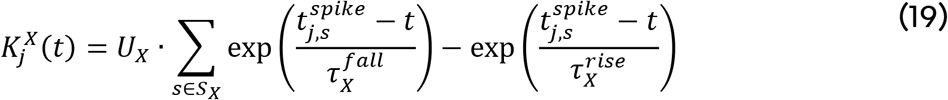

is the postsynaptic potential of a neuron *j*, where ***X*** represents either excitatory or inhibitory neurons (E or I), 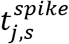 is the time of a spike *s* ∈ *S_X_* where *S_X_* denotes all spikes emitted up to time *t*, 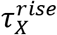 and 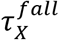 are the time constants of the rise and fall of postsynaptic potential, with amplitude factors *U_E_*=0.4 nA and *U_I_*=0.6 nA. We set 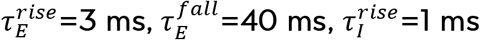, and 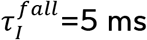.

Spikes were triggered when the membrane potential (equation (17)) of a neuron reached −15 mV from a value below the threshold. At that time, the potential was set to 100 mV for 1 ms, then reset to −65 mV for a 3 ms absolute refractory period. Connection weights 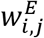 and 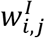 were set such that 30% of neurons were inhibitory.

The out-degree connectivity was set to 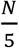 for individual E cells and 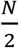 for I cells. Connection weights were structured at random following a uniform distribution with 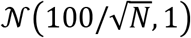 while ensuring that all outgoing connections of E cells were positive and all outgoing connections of I cells were negative, in line with Dale's law. Simulations employed a forward Euler method with a time resolution of 0.1 ms. Null and potent modes of **W**_0_ were set according to equation (14) after replacing the activity of the mean-rate model (**X**) with temporally smoothed spikes (100 ms rolling window).

### 4.3 Experimental data

Data analysed in this study were obtained from a public repository (crcns.org) and described in detail in related work [16]. These data were composed of simultaneous extracellular recordings from neuronal populations in output layers (2/3-4B) of V1 as well as their primary target in middle layers of V2. Recordings were performed in 3 *macaca fascicularis* under sufentanil anesthesia using a Utah array (V1) and tetrodes (V2). During these recordings, subjects were shown oriented gratings for a brief duration (1.28 sec) followed by a blank screen (1.5 sec). A range of 88-159 V1 neurons (mean: 112.8) and 24-37 V2 neurons (mean: 29.4), including both well-isolated single units and multi-unit clusters, were analysed. A total of 400 trials were examined for each of 8 stimulus orientations. The receptive fields of V1 and V2 neurons were aligned retinotopically, thus promoting feedforward interactions.

All data analyses were performed using custom scripts in the Matlab language (MathWorks, Natick MA). Statistical analyses were performed with a one-sided non-parametric Wilcoxon rank sum test for non-normal data.

### 4.4 Ridge regression

Ridge regression aimed to minimize a loss function [50],

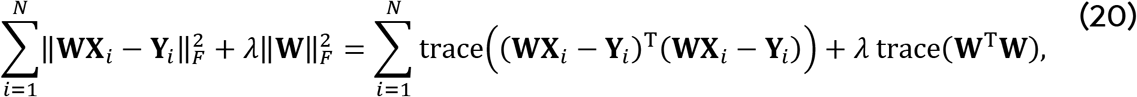

where **X**_*i*_ is the activity of *i* ∈ *N* sender neurons, **Y**_*i*_ is the activity of receiver neurons, **W** are regression weights, and *λ* is a regularization term. By expansion,

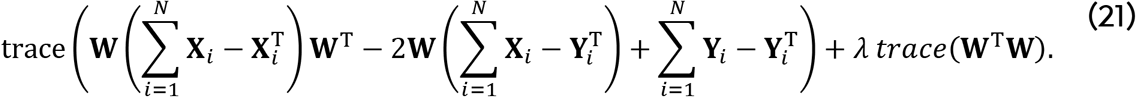

Taking the derivative with respect to **W**,

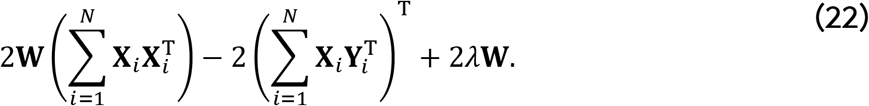

Setting the derivative to zero and solving yielded

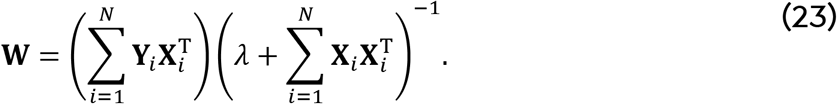

The least-squares approximation of receiver activity was obtained by 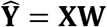. The regularization term *λ* was chosen such that the loss function (equation (20)) was no more than 5% lower than the corresponding unregularized (*λ* =0) value on a given trial.

## Acknowledgments

This work was supported by a Discovery grant to J.P.T. from the Natural Sciences and Engineering Council of Canada (NSERC Grant No. 210977).

## Author contributions

J.P.T. and A.P. designed and conducted the experiments, and analyzed the results. Both authors wrote and reviewed the manuscript.

## Data availability

Custom Matlab code is available from the corresponding author upon request.

